# Characterization of antioxidant properties and metabolite profile of *Agave atrovirens* extracts

**DOI:** 10.1101/184226

**Authors:** Victor Olvera-Garcia, Ana Belen Granado-Serrano, Mariona Jové, Anna Cassanyé, Anaberta Cardador-Martínez, Manuel Portero-Otin, José CE Serrano

## Abstract

*Agave spp*. is widely analyzed because several functional properties have been described. Some minor bioactive compounds such as polyphenols, saponins and Maillard compounds produced during extraction procedures have been reported to exert antioxidant properties. The objective of this study was to elucidate the antioxidant properties of three different *Agave atrovirens* extracts in a HepG2 cell culture assay. The three extracts analyzed mostly showed antioxidant properties with an increase in NRF2 content in nuclear extracts. However, a differential response was observed in the reduction of protein oxidative damage in the three extracts analyzed, the crude extract being the one that mainly induced a reduction in oxidative damage. Metabolomic analysis was performed to elucidate the potential molecules responsible for the antioxidant properties, where 2-amino-4-methylphenol could be the main candidate responsible for inducing the transcription of cellular antioxidant response elements. It could be concluded that crude extract of *Agave atrovirens* may increase the cellular antioxidant defense system, with a reduction in oxidative damage

**List of abbreviations**
AAPH: 2,2´-azobis(2-amidinopropane) dihydrochloride
ABTS: 2,2'-azino-bis(3-ethylbenzothiazoline-6-sulphonic acid)
AGEs: Advanced glycated end-products
ARE: Antioxidant Response Elements
DNP: Dinitrophenyl
EB: *Agave atrovirens* butanolic extract
EC: *Agave atrovirens* crude extract
ET: *Agave atrovirens* total extract
FRAP: ferric reducing antioxidant power
MDA-lys: malondialdehyde-lysine
NRF2: Nuclear factor (erythroid-derived 2)-like 2
ORAC: oxygen radical antioxidant capacity
PARP: Poly ADP ribose polymerase
TE: Trolox equivalents
TPTZ: 2,4,6-tri(2-piridil)-s-triazina

## Introduction

Since ancient times agaves have been used in Mexico as source of food, drink, medicine, fuel, shelter, ornament and fibers. More recently, just few species are exploited and the expansion of commercial monocultures for industrial purposes has threaten their biodiversity and generated ecological problems like soil erosion [1]. The empirical medicinal use as well as the supposed beneficial health effects like the low glycemic index of the agave fructose-rich syrups [2,3] has increased the interest in studying species that are not used traditionally like *A. atrovirens*. Additionally, documented productivity for species like *A. mapisaga* or *A. salmiana* suggests that non-commercial exploited species could may have equal or better productivity than *A. tequilana*, the most cultivated specie [4].

Recently, a significant *in vivo* reduction in malondialdehyde content in serum of rats fed with *Agave tequilana* has been reported. Similarly, a reduction in protein oxidative damage was observed in the cecum [5], suggesting the presence of antioxidant compounds in this plant genus. Many species of the genus *Agave* constitute an important source of steroidal sapogenins, mainly hecogenin. The amounts of total phenol content and total flavonoid content may range from 10.5 to 39.3 gallic acid equivalents (mg/100 g of dry plant matter) and from 43.3 to 304.8 catechin equivalents (mg/100 g of dry plant matter), respectively [6].

Despite antioxidant capacity have been reported for some Agavacea members like *A. utahensis, A. sisalana* and *A. furcroydes* [7-9], the antioxidant effect has been demonstrated only in the traditional chemical assays and it is not clear if these antioxidant properties are maintained in a living cell or how it is related to the cytotoxic effect. In this work we evaluated the effect of the *A. atrovirens* extracts on the activation of the antioxidant defense system via the traslocation of NRF2 in HepG2 cells as well as its effects in the oxidative damage of proteins. A chemical identification was done in order to establish a possible relation between the chemical composition of the extracts and their biological activity.

## Materials and Methods

### *Agave atrovirens* extracts

*Agave atrovirens* aqueous extracts were kindly provided by the company Nutracéuticos de Agave S.P.R de R.L. Crude extract (EC) was obtained by grinding the fresh leaves of *A. atrovirens*, posteriorly, it was processed until a dark-brown liquid, free of waxes and a solid content of 20 º Brix was obtained (Total extract, ET). Both extracts were lyophilized and stored at −40 ºC until used. Butanolic extract (EB) was obtained by dissolving 20 g of ET in 80 mL of H_2_O and extracted with 300 mL of n-Butanol. The organic phase was evaporated until dryness under reduced pressure using a rotavapor system.

### Antioxidant capacity assays

Antioxidant capacity assays performed in extract and cell lysates followed the procedure described for ORAC, FRAP and ABTS assays [11–13]. Results were expressed as micromoles of Trolox equivalents.

### Cell culture & treatments

Human HepG2 cells were grown in a humified incubator containing 5% CO_2_ and 95% air at 37ºC. Cells were grown in DMEM F-12 medium suplemented with 10 % of FBS, 100 UI/mL of Penicillin and 100 µg/mL of Streptomicyn. In all experiments, cells were grown until confluence and then transferred to 6 or 96-well plates. Stock solutions (10 mg/mL for EC and ET) and (1.0 mg/mL for EB) were prepared dissolving in DMSO each extract. These solutions were diluted in serum-free medium at 12.5, 25.0, 50 µg/mL for EC and ET, and 1.0, 2.0, 3.0 µg/mL for EB. The concentrations of the extracts were determined after cell viability assays, in the case of EB extract below 3.0 µg/mL no toxicity was observed. The day of the assay the medium was changed to serum free medium and cells were exposed to the extracts or during 4 h for oxidative damage or 6 h for NRF2 analysis.

### Preparation of total lysates for protein oxidative stress markers analysis

Oxidative lesions were evaluated using anti-DNP (Dinitrophenyl) for carbonyl adducts formation (Sigma D9565), malondialdehyde-lysine (MDA-Lys) adducts (Abcam ab20703), AGEs (advanced glycation end-products) (Transgenic Inc KAL-KH025-02) and degree of protein ubiquitination (Sigma U5379) as markers. After treatments, HepG2 cells were washed with cold PBS and lysed at 4 ºC using RIPA buffer with protease and phosphatase inhibitors. Protein concentrations in the lysates were measured using the Bradford assay with bovine serum albumin as a standard. Samples were stored at −80 ºC until used for Western blot analysis.

### Preparation of nuclear and cytosolic extracts for NRF2 analysis

To analyze NRF2 (Santa Cruz Biotechnology sc13032), cells were resuspended at 4 ºC in Buffer A (10 mM HEPES pH 7.9, 10 mM KCl, 0.5 mM dithiothreitol (DTT), 0.2 mM phenylmethylsulfonyl fluoride (PMSF) and 1.5 mM MgCl_2_), allowed to swell on ice for 10 min, and then vortexed for 10 s. Samples were centrifuged at 10,000g for 2 min and the supernatant containing the cytosolic fraction was stored at −80 ºC. The pellet was suspended in cold buffer B (20 mM HEPES pH 7.9, 25% glycerol, 420 mM NaCl, 1.5 mM MgCl_2_, 0.2 mM EDTA, 0.5 mM DTT, 0.2 mM PMSF, 2.5 µg/mL leupeptin and 2.5 µg/mL aprotinin) and incubated on ice for 20 min for high salt extraction. Cellular debris was removed by centrifugation at 13,000g for 10 min at 4 ºC. Supernatants fractions containing cytosolic or nuclear protein extracts were analyzed for protein concentration as described above. Both cell extracts were stored at −80 ºC until use for Western blotting.

### Western blot analysis

Equal amounts of proteins were resolved by SDS-polyacrylamide gel electrophoresis (SDS-PAGE) and electrobloted onto polyvinylidene difluoride membranes (Immobilon-P Millipore, Bedford, MA, USA). Immunodetection was performed using the corresponding primary antibody followed by incubation with peroxide-conjugated anti-rabbit (Pierce Biotechnology 31450) or anti-mouse (GE Healthcare NA931V) immunoglobulin. PARP (Poly ADP ribose polymerase) (Santa Cruz Biotechnology sc7150) was used as a marker for the nuclear extract. For total lysates a monoclonal antibody to β-actin (Sigma A5441) was used to control protein loading. Normalization of AGEs, DNP, MDA-Lys, ubiquitin and NRF2 western blots were ensured by staining the membranes with the Gallyas method. Protein bands were visualized by the chemiluminescent ECL method (Millipore Corporation, Billerica, MA, USA) and quantified in a ChemiDoc MP imaging system (Bio-Rad Laboratories Inc., CA, USA) using the Image Lab software Version 4.0.

### Metabolomics analysis

Metabolites were extracted from EC, ET or EB with a mixture of water/methanol and BHT 1mM as antioxidant. After mixing in a vortex, samples were incubated 1 hr at −20 ºC and centrifuged at 13000 rpm/4 ºC/30 min. Supernatants were evaporated until dryness using a Speed Vac system and dissolved with 50 µL of methanol and 50 µL of aqueous acetic acid 0.4%. Samples were filtered and injected to an Agilent 1290 LC system (G2226A model) coupled to an ESI-Q-TOF MS/MS 6520 instrument (Agilent Technologies, Barcelona, Spain). 2 μL of extracted sample was applied onto a reversed-phase column (Zorbax SB-Aq 1.8 µm 2.1 × 50 mm; Agilent Technologies) equipped with a precolumn (Zorba-SB-C8 Rapid Resolution Cartridge 2.1 × 30 mm 3.5 µm; Agilent Technologies) with a column temperature of 60 °C. The flow rate was 0.6 mL/min. Solvent A was composed of water containing 0.2% acetic acid and solvent B was composed of methanol 0.2% acetic acid. The gradient started in 2% B and increased to 98% B in 13 min and hold at 98% B for 6 min. Data were collected in positive and negative electrospray mode TOF operated in full-scan mode at 100-3000 m/z in an extended dynamic range (2 GHz), using N_2_ as the nebulizer gas (5 L/min, 350 ºC). The capillary voltage was 3500V with a scan rate of 1 scan/s. The Masshunter Data Analysis Software (Agilent Technologies, Barcelona, Spain) was used to collect the results, and the Masshunter Qualitative Analysis Software (Agilent Technologies, Barcelona, Spain) was used to obtain the molecular features of the samples using the Molecular Feature Extractor (MFE) algorithm (Agilent Technologies, Barcelona, Spain). Finally, the Masshunter Mass Profiler Professional Software (Agilent Technologies, Barcelona, Spain) was used to perform a non-targeted metabolomics analysis of the extracted features.

### Statistical analysis

Statistical analyses for metabolomics analysis were done using the Mass Profiler Professional Software (Agilent Technologies, Barcelona Spain). Normal distribution of the variables was checked by the Kolmogorov-Smirnoff test.

Correlation between molecular features was evaluated by the Pearson correlation coefficient. Otherwise, statistics calculations were performed using the computer software Statistica (8.0 Stat Soft, Inc. Tulsa OK, USA). Data were subjected to ANOVA procedures and were significant differences existed; means were separated by the Tukey test (P<0.05).

## Results and Discussion

### Antioxidant capacity

The chemical composition of the extracts may be differentiated in terms of the solubility of its components. For instance, the crude extract (EC) contains hydrophilic and a lipophylic compounds, while the butanolic extract (EB) is rich in lipophylic compounds, mainly saponins. Antioxidant capacity was determined in terms of its equivalency with Trolox, a water-soluble vitamin E analog and measured by the FRAP and ORAC assay (Table 1). Differences in antioxidant capacity could be expected within methods and may respond differentially to the antioxidant profile in the extract. For instance, the phenolic content of EC is 62% higher than the ET as well as a 222% higher in acidity properties [14], which may explain the increased antioxidant capacity observed in EC by the FRAP assay. While the high hydroxymethylfurfural content in EC [14], due to its high reactivity may reduce the reported value of antioxidant capacity of EC in the ORAC assay [13,15]. The content of other antioxidants like ascorbic acid have been reported also in the literature [16] that could explain part of the antioxidant capacity observed.

**Table 1.**
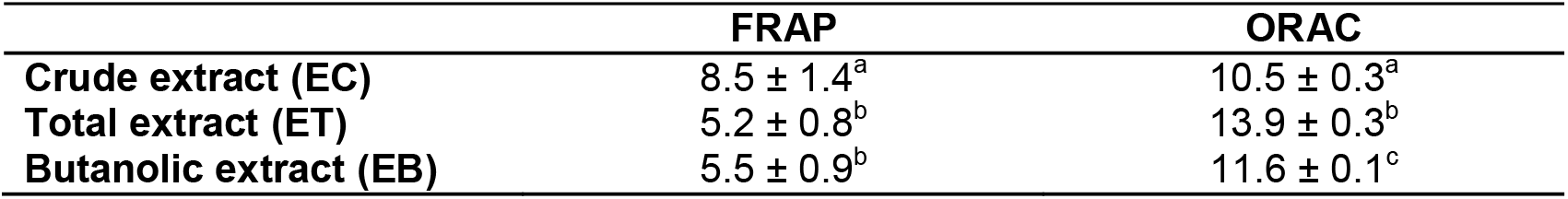
Antioxidant capacity of *Agave atrovirens* extracts measured by the FRAP and ORAC assay. Results are expressed as μmol of Trolox equivalents per g of freeze-dried extract. Different letters within columns express statistically differences between samples (p<0.05). Differences were determined by one-way AnOvA and Tuckey post-hoc analysis.

|  | FRAP | ORAC |
| --- | --- | --- |
| **Crude extract (EC)** | $8.5 \pm 1.4^a$ | $10.5 \pm 0.3^a$ |
| **Total extract (ET)** | $5.2 \pm 0.8^b$ | $13.9 \pm 0.3^b$ |
| **Butanolic extract (EB)** | $5.5 \pm 0.9^b$ | $11.6 \pm 0.1^c$ |

### Antioxidant properties

Antioxidant properties were measured in terms of the quantification of protein oxidative damage in HepG2 cell culture after the exposure of 4 h of Agave extracts, and by the determination of NRF2 translocation to the nucleus after 6 h of exposure. Several plant foods have reported the ability to increase the expression of NRF2 and related antioxidant enzymes [17] that make it a suitable marker of antioxidant properties. In this context, a high antioxidant property was defined as a reduced protein oxidative damage together with an increased translocation of NRF2 to the nucleus. To simplify the interpretation of the described antioxidant properties, a graph with quadrants was used as shown in Figure 1, where Q1 represents Antioxidant Property (low protein oxidation and high NRF2 translocation), Q2 represents Antioxidant Capacity (low protein oxidation and low NRF2 translocation), Q3 represents oxidative properties together with cell detoxification system response activation (high protein oxidation and high NRF2 translocation), and Q4 represent oxidative properties without cell antioxidant response (high protein oxidation and low NRF2 translocation). In this context, a desirable product with antioxidant properties should be able to reduce protein oxidation as well as an induction of the cell antioxidant defense (Q1); or at least reduce the protein oxidative damage (Q2).

**Figure 1.**
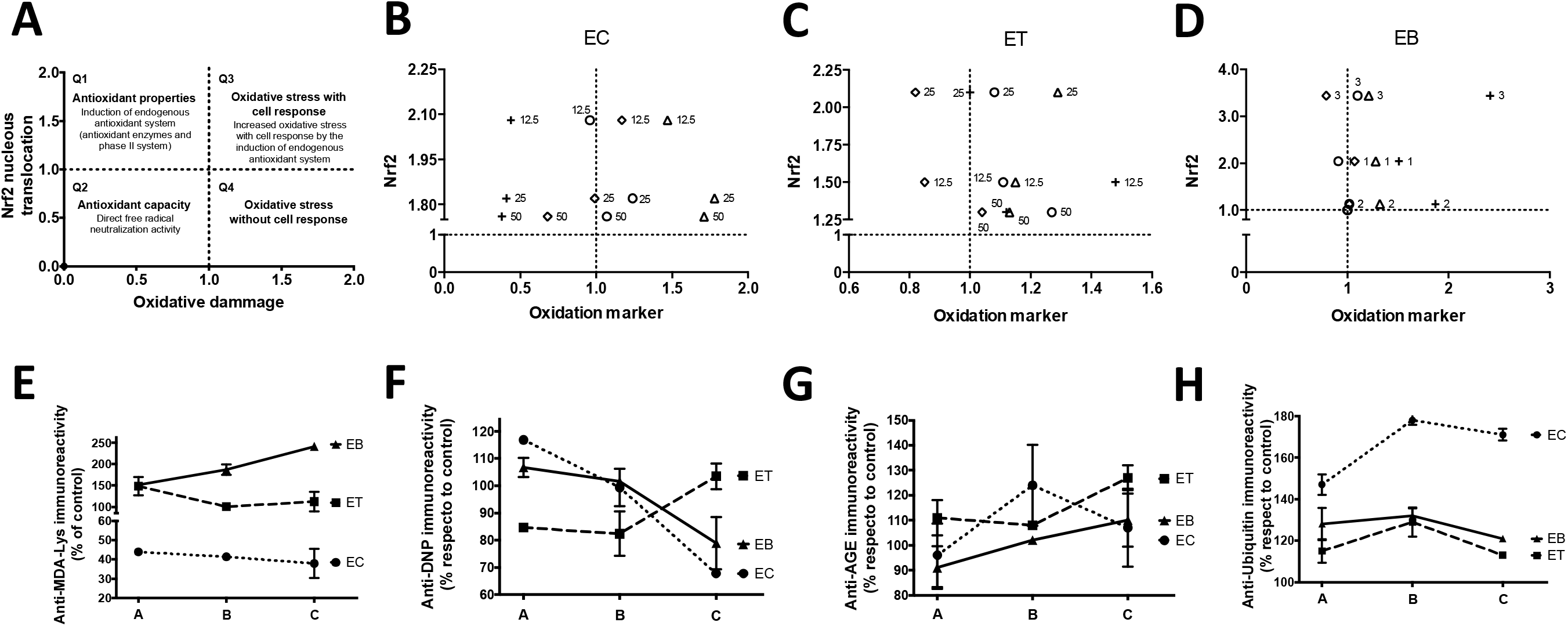
Antioxidant capacity and properties of Agave extracts. **A** Shows the general scheme of evaluation of the antioxidant property (activation of Nrf2 translocation to the nucleus), antioxidant capacity (reduction in oxidative markers) and oxidative stress (an increase in oxidative marker compared to a control condition). **B, C** & **D** shows the antioxidant properties and capacity evaluation of the three extracts analyzed Crude extract (CE), Total extract (TE) and Butanolic extract (BE). (+), (◊), (o) and (Δ) express the immunoreactivity to MDA-lys, derivitized carbonyls with DNP, advanced glycation end-products and ubiquitin respectively as oxidation markers. From **E** to **H** the dose response effect of the three extracts within the different oxidation markers. The A to C concentrations represents the doses of 12.5, 25 and 50 μg/mL respectively in EC and ET, and 1, 2 and 3 μg/mL in EB. Statistical significant correlation was observed in the increase of MDA-lys oxidation by EB (p=0,0012 r^2^=0,9436) and the reduction in carbonyl adducts (DNP) by EC (p=0,0006).

All agave extracts analyzed showed a capacity to induce the translocation of NRF2 to the nucleus (Figure 1 b-d), being the EB extract the one that showed the higher ratios. It could also be observed that the rate of NRF2 translocation is only dose-dependent in the EC extracts, while the ET and EB extracts showed U shape responsiveness. In this sense, it must be taken into account that the activation of NRF2 could be due, in one hand to increased oxidative stress or otherwise because of the extract effects on the NRF2 activation without an increase in oxidative stress. The latter scenario is the most desirable between both, since this would imply that the agave extract does not induce oxidative damage. In general terms, it could be described that the EB extracts induction of NRF2 activation could be attributed to the increase in oxidative stress (Figure 1d), while the EC extract showed antioxidant properties (Figure 1b). Moreover, it could be suggested that at low doses (12.5 μg/mL) of the EC extract, the reduction in oxidative stress could be attributed to the activation of NRF2 pathway, while at higher doses (50 μg/mL) the observed reduction could be ascribed to the antioxidant capacity content of the extract.

To determine if the increase in NRF2 activation was in response to an increase in oxidative stress, protein oxidative damage was determined as shown in Figure 1 e-h. Respect to control cells, only the EC extract was able to reduce the amount of protein oxidation measured by MDA-lys and DNP. Both EB and ET extracts induced a higher protein oxidation rate even at low doses. In this respect, the increase in the concentration of the EB extract induced a higher oxidation rate of MDA-lys compared to control cells. Protein ubiquitination was considerable higher respect to control cells in the EC treated cells (Figure 1h). This higher ubiquitination could explain in part the low grade of protein oxidation observed in the EC treated cells since it may denote a higher protein turnover. The increase in protein ubiquitination could be explained as well by the increase in the activation of NRF2 observed also in cells exposed to EC, since NRF2 may directly active ubiquitin/proteosome genes [18]. Therefore, it could be suggested at this point that the lower degree of oxidative stress observed in the EC treated cells could be associated with the increase in activation of the detoxification system by NRF2 as well as the induction of an increase in protein turnover.

In order to verify the increase in the antioxidant properties of the agave extracts, the antioxidant capacity of the cells lysates was measured (Figure 2a). Compare to control cells and to cells treated by the ET and EB extracts, EC extract was the only one able to induce changes in antioxidant capacity content of the cells lysates. Moreover, the observed changes were in an inverse dose-dependent manner and similarly as the NRF2 nucleus content (Figure 1b). Suggesting, that NRF2 could be related to the increase in the cytosolic antioxidant capacity due to an increase in the antioxidant response elements activation. To confirm this hypothesis, a kinetic measurement of the FRAP assay was developed based on the premise that antioxidant enzyme activity will induce a continuous increase in absorbance in the FRAP reaction mixture. Since most non-enzymatic antioxidants induce a rapidly increase and later stable measure in absorbance in the FRAP assay, e.g. the slope for the Trolox calibration curve was between 0.03 and 0.05. In this sense, the EC 12.5 treated cells showed a significant increase ability of continuously induce antioxidant capacity over the time, which may suggest a higher antioxidant enzymatic defense.

**Figure 2.**
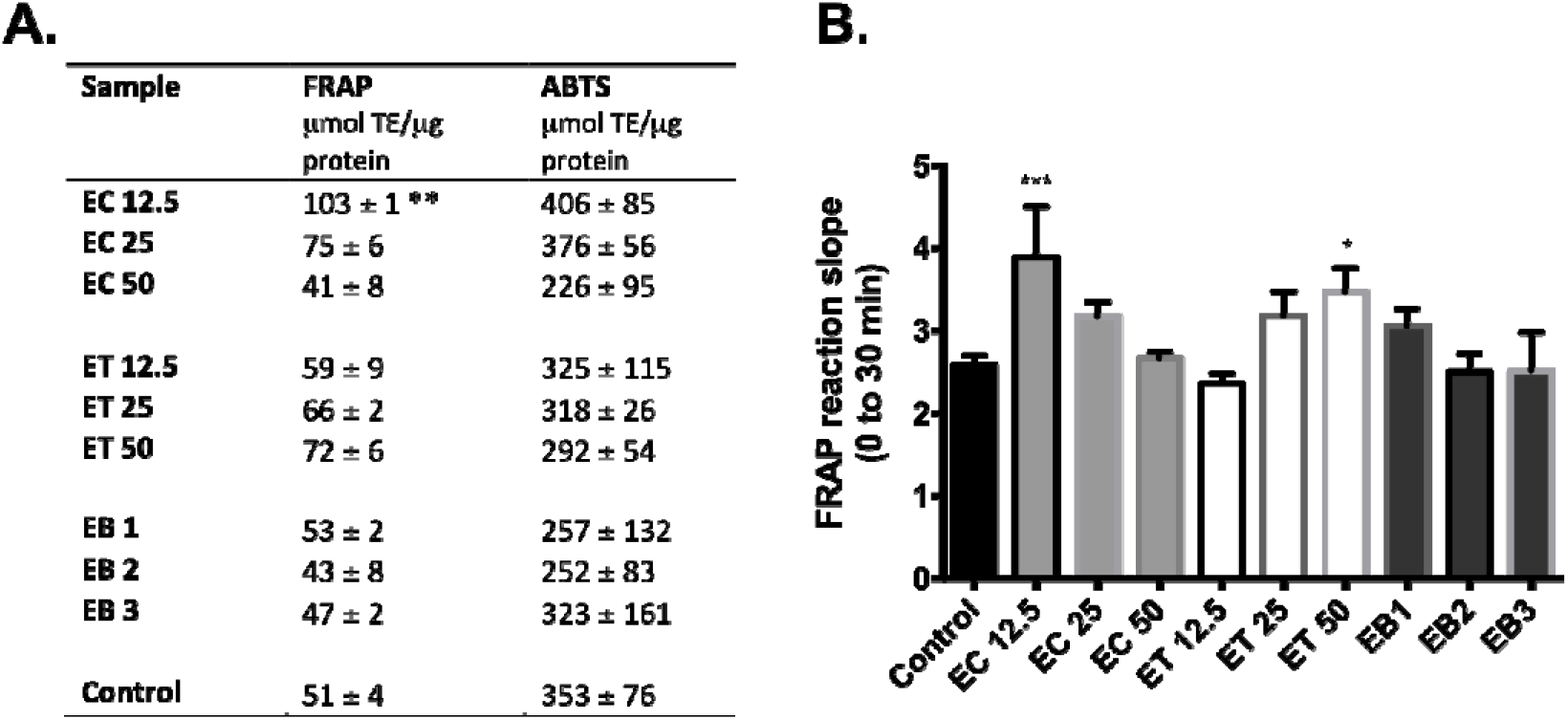
Antioxidant capacity of cell extracts measured by FRAP and ABTS assays. **A.** HepG2 cytosolic antioxidant capacity after the exposure of agave extracts at different concentrations during 4 h. Significant increase in antioxidant capacity respect to control was observed in cells exposed to EC 12.5 measured by FRAP (p=0.0056). **B.** FRAP kinetic antioxidant capacity measurement. The increase in absorbance (λ 595 nm) from 0 to 30 min was registered; the figure denotes the slope of the absorbance per time (minutes). Significant higher slope respect to control was observed in the samples EC 12.5 (p=0.0005) and ET 50 (p=0.0140). The slope of trolox standards was between 0.03 and 0.05, which may denote a complete reaction at minute 0.

It could be concluded from this part, that agave extracts *in vitro* antioxidant capacity measured both by FRAP and ORAC is comparably lower than most edible plants. However, the most important finding of this study was the evidencing of the antioxidant properties via the activation of NRF2 and possibly the antioxidant response element (ARE) signaling pathway. Which may be responsible of the reduction in the protein oxidative damage observed in the HepG2 cell culture. No reports have been found in the literature that describes this effect from Agave genus.

Notwithstanding, the antioxidant properties together with and low protein oxidative damage was observed solely in the crude hydrophilic extract. Suggesting that the further concentration and extraction procedures employed, mainly in the total and butanolic extracts, removes or inhibits the action of the bioactive compounds responsible of the observed effects in the crude extract.

### Metabolomic analysis of Agave extracts

The chemical composition of the three extracts is considerable different as observed in the heat map and PCA analysis of the metabolites (Figure 3). The chemical composition of the EC extract shows more similarities to the ET extract (3369 common metabolites) compared to EB extract (1112 common metabolites). These similarities are also observed in the antioxidant properties described before (Figure 1), where the antioxidant behavior of the ET extract resulted to be in the middle of the observed effects of EC and EB extracts. At least 319 metabolites resulted to be differentially present in the EC extract and not in the ET and EB extracts. From these, 33 could be identified by the MassHunter Agilent Technologies PCDL database (**Supplementary Table S1**, grey lines). To determine which of these compounds could be responsible of the antioxidant properties a review of compounds bioactivity was performed in PubChem database (www.pubchem.ncbi.nlm.nih.gov), from which, 5 compounds were found to report bioactivity related to antioxidant defense and inflammatory pathways (**Supplementary Table S2**). Although it is possible that the observed effect could be attributed to the synergism between compounds, the 2-amino-p-cresol has been reported to be an agonist of the antioxidant response elements by a direct interaction with NRF2 protein. Similarly, (±)-Taxifolin could act as inhibitors of FAD-linked sulfhydryl oxidase, which could be related to a reduction in the cellular oxidative stress. The other metabolites identified are inhibitors of inflammatory responses that are known as possible generators of free radicals and oxidative stress.

**Figure 3.**
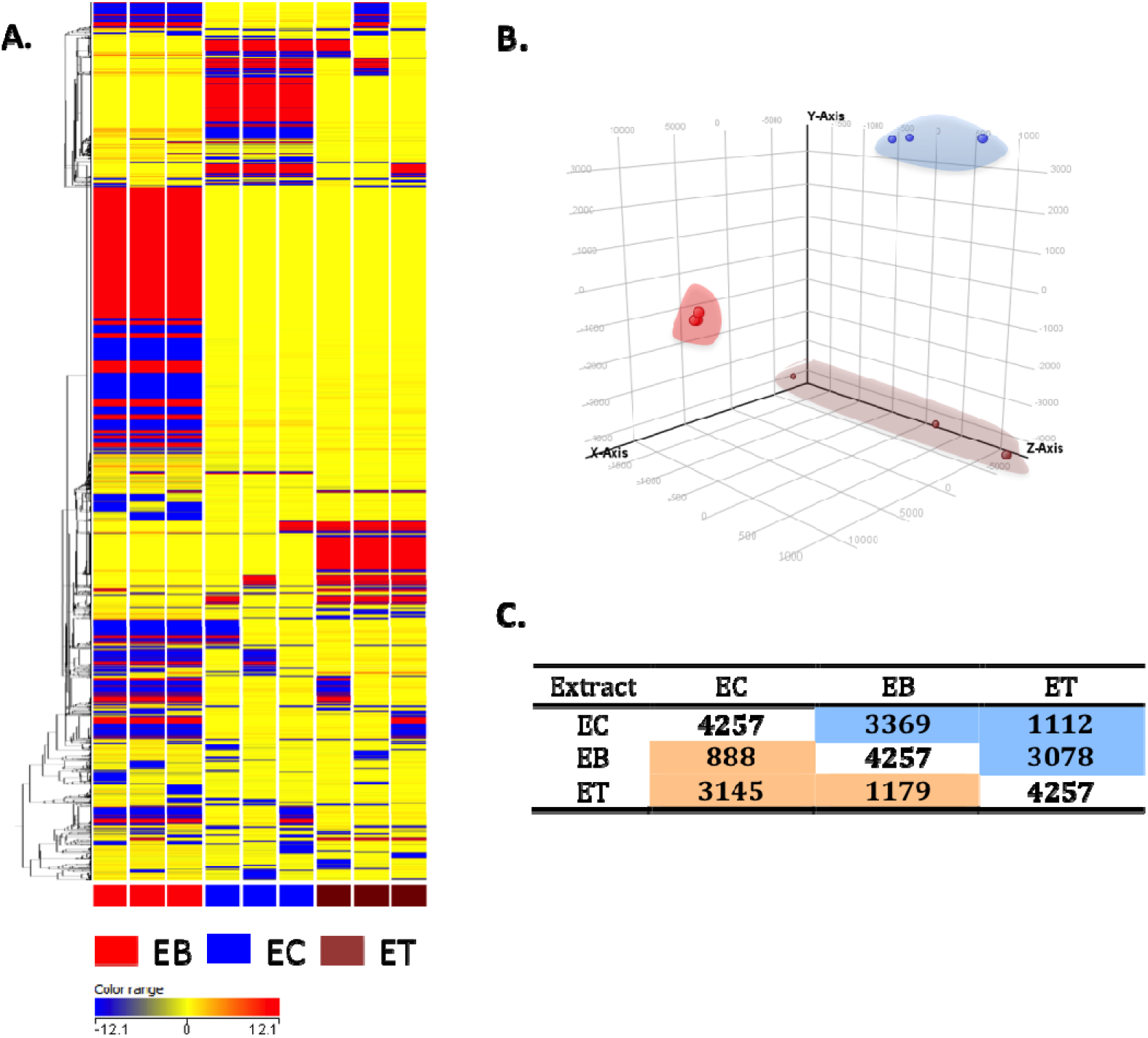
Metabolomic analysis differences between the agave extracts. **A**. Heat map showing the molecular features found in the agave extracts. Scale map for heat intensity is shown below. **B**. PCA graph demonstrating the differentiation of agave extracts metabolomic profiles. Brown spots represent EC; blue spots ET and red spots EB. C. Entities found to be differentially expressed are represented in the blue box, while entities found not to be differentially expressed are represented in the orange box.

Although 5 compounds could be identified and related to the activation of NRF2, other low and high molecular weight compounds (under 100 and above 3000 kDa) and lipophylic compounds (terpenoids) not included in the chromatographic analysis and possibly contained in the extracts could also be agonist of NRF2 signaling pathway. For example, Maillard reaction products, with a high molecular weight, have been reported to have the ability to activate NRF2 transcription factors in macrophages and Caco-2 cells [19]. Additionally, several authors have described that saponins from different sources are also NRF2 activators [20-22] and Agave plants are important sources of saponins and the occurrence of steroidal saponins is well documented [23,24]. On the other hand, it is worthy to mention that cytotoxic properties have been described for agave saponins [7,25]. This fact could explain the elevated oxidative damage observed in the butanolic extract, which are supposed to be rich in saponins.

It could be concluded that the crude extract of *Agave atrovirens* seems to possess antioxidant properties via the activation of NRF2 antioxidant response elements transcription. Further studies should be developed to confirm the bioactivity of the recognized metabolites and the technological procedure to increase its content in the crude extract.

## Acknowledgment

The study was performed thanks to the financial support of the Consejo Nacional de Ciencia y Tecnología (CONACyT) México and to the doctoral scholarship number 310208. This research article has received a grant for its linguistic revision from the Language Institute of the University of Lleida (2015 call)

## Conflict of interest

The authors declare to have no potential conflict of interest.

